# Species tree disequilibrium positively misleads models of gene family evolution

**DOI:** 10.1101/2020.01.08.899518

**Authors:** M. Elise Lauterbur, Sarah Heder, Laurel R. Yohe, Liliana M. Dávalos

## Abstract

Gene duplication is a key source of evolutionary innovation, and multigene families evolve in a birth-death process, continuously duplicating and pseudogenizing through time. To empirically test hypotheses about adaptive expansion and contraction of multigene families across species, models infer gene gain and loss in light of speciation events and these inferred gene family expansions may lead to interpretations of adaptations in particular lineages. While the relative abundance of a gene subfamily in the subgenome may reflect its functional importance, tests based on this expectation can be confounded by the complex relationship between the birth-death process of gene subfamily evolution and the species phylogeny. Using simulations, we confirmed tree heterogeneity as a confounding factor in inferring multi-gene adaptation, causing spurious associations between shifts in birth-death rate and lineages with higher branching rates. We then used the *olfactory receptor* (*OR*) repertoire, the largest gene family in the mammalian genome, of different bat species with divergent diets to test whether expansions in olfactory receptors are associated with shifts to frugivorous diets. After accounting for tree heterogeneity, we robustly inferred that certain *OR* subfamilies exhibited expansions associated with dietary shifts to frugivory. Taken together, these results suggest ecological correlates of individual *OR* gene subfamilies can be identified, setting the stage for detailed inquiry into within-subfamily functional differences.

## Introduction

Gene duplication is a key source of evolutionary innovation, facilitating the evolution of new functions of duplicated genes, and even the acquisition of new and specific biological roles (Assis and Bachtrog 2013). In contrast with most protein-coding regions of the genome for which duplication is relatively rare, certain regions duplicate frequently (Nei 1969) and, released from purifying selection, mutation pseudogenizes some of the copies over time (Nei and Hughes 1991) generating multigene families. Unlike single copy genes, multigene families evolve through a process of duplication (birth) and pseudogenization and deletion (death). Assuming multiple copies as a starting point, an equilibrium between births and deaths will result in turnover in the composition and identity of the genes (Han et al. 2013), independent of gene identity and function. Since multigene families evolving through the birth-death process encode for important functions such as chemosensation (Nei et al. 1997), immune defense (Nei et al. 1997), and even development (Nei and Rooney 2005), their evolutionary rates and shifts in duplication or loss rates are of broad interest in molecular evolution, particularly when shifts correspond to major ecological transitions.

There are two main approaches to infer changes in duplication and loss: (1) estimating the rates of birth and death along branches of the species tree inferred from the number of gene copies present in each species and (2) and gene tree-species tree reconciliation to infer incongruences that suggest a duplication or loss event (Yohe et al. 2019). In the former approach, multi-gene family evolution models typically parameterize a probability distribution over the counts of the number of genes within the family from the tips of the species tree (Bie et al. 2006). This parameterization thus integrates the birth-death model and the structure of the phylogeny, making it possible to test for equilibrium in the birth-death model for a gene family, as well as pinpointing the specific branch on the tree responsible for disequilibrium, if found. In similar integrated models, however, disequilibrium can arise from heterogeneity in the birth-death process (the intended hypothesis test), in the branching process of the phylogeny (not intended), or both, thus confounding tree and gene subfamily disequilibria. First identified in binary state-dependent speciation (-SSE) models (Rabosky and Goldberg 2015), this problem has been found to be more general (Rojas et al. 2018) when there is heterogeneity in the branching process of the phylogeny (Beaulieu and O’Meara 2016). In other words, a rejection of the null emerges not from the evolution of the trait of interest, but from rejecting the embedded assumption that a single lineage branching rate applies to the entire phylogeny when it does not (Beaulieu and O’Meara 2016). Although hitherto unexplored for inference of multi-gene family evolution, previous findings for -SSE models suggest caution in applying hypothesis tests using birth-death models to phylogenies with known heterogeneity in their branching pattern.

To determine how species tree heterogeneity affects birth-death models for multigene families, particularly when linked to ecological diversity, three components are needed: (1) a diverse multigene family across species with divergent ecologies; (2) shifts in ecological function that may relate to gene family evolution; and (3) a system with strong shifts in diversification rates that may lead to tree heterogeneity. Olfactory receptor (*OR*) proteins are encoded by precisely such a diverse gene family, comprising the largest protein-coding fraction of a given mammalian genome. Although rich OR repertoires have been recorded across vertebrates (Vandewege et al. 2016), the greatest OR diversity is found in mammals, with hundreds of *olfactory receptor* (*OR*) genes encoded in tandem arrays of similar copies (Niimura and Nei 2003). If mammalian ecology changes such that the sense of smell is no longer usable, as among many aquatic mammals, disequilibrium ensues with higher rates of loss. In an extreme case, odontocete whales — dolphins, porpoises, and sperm whales among others — lack an olfactory bulb, as well as other morphological and molecular correlates of olfaction, and a high proportion of their olfactory subgenome is pseudogenized (74-100%) compared to terrestrial relatives (McGowen et al. 2014). In contrast, when function is important, selection will act against the random pseudogenization or deletion of copies, resulting in neofunctionalization of the duplicated gene or the retention of more functional gene copies and evidenced by expansion of specific gene subfamilies. As a result, comparing the relative sizes of gene subfamilies across species and inferring the corresponding duplications and losses can yield evidence for selection on particular gene subfamilies, or shifts in selection therein (figure 1).

**Figure 1.**
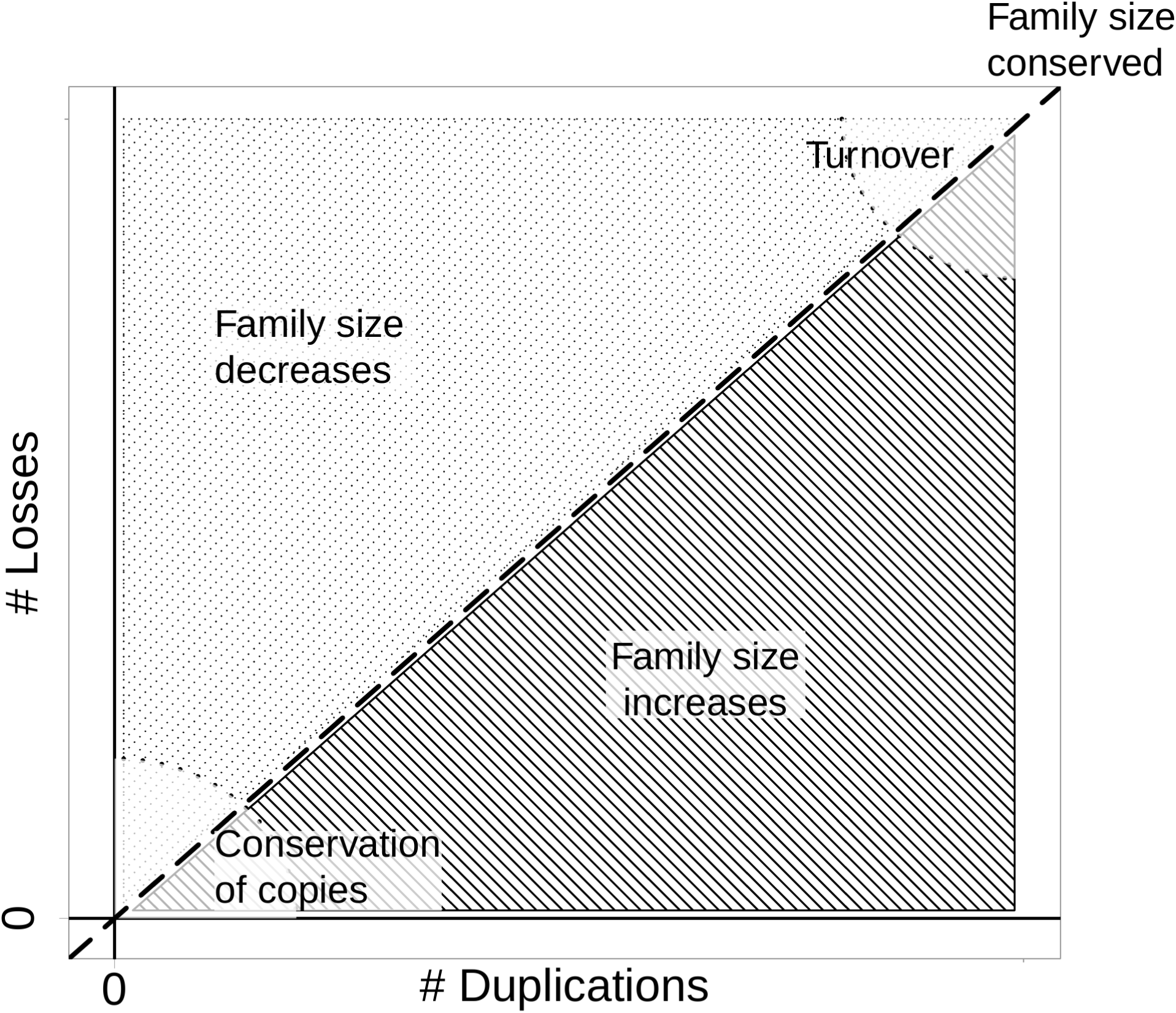
Relationship between the duplications, losses and multi-gene family size. The dashed line indicates equilibrium between losses and duplications, turnover indicates changes in the identity of genes over time resulting from high duplication and loss rates.

**Figure 2.**
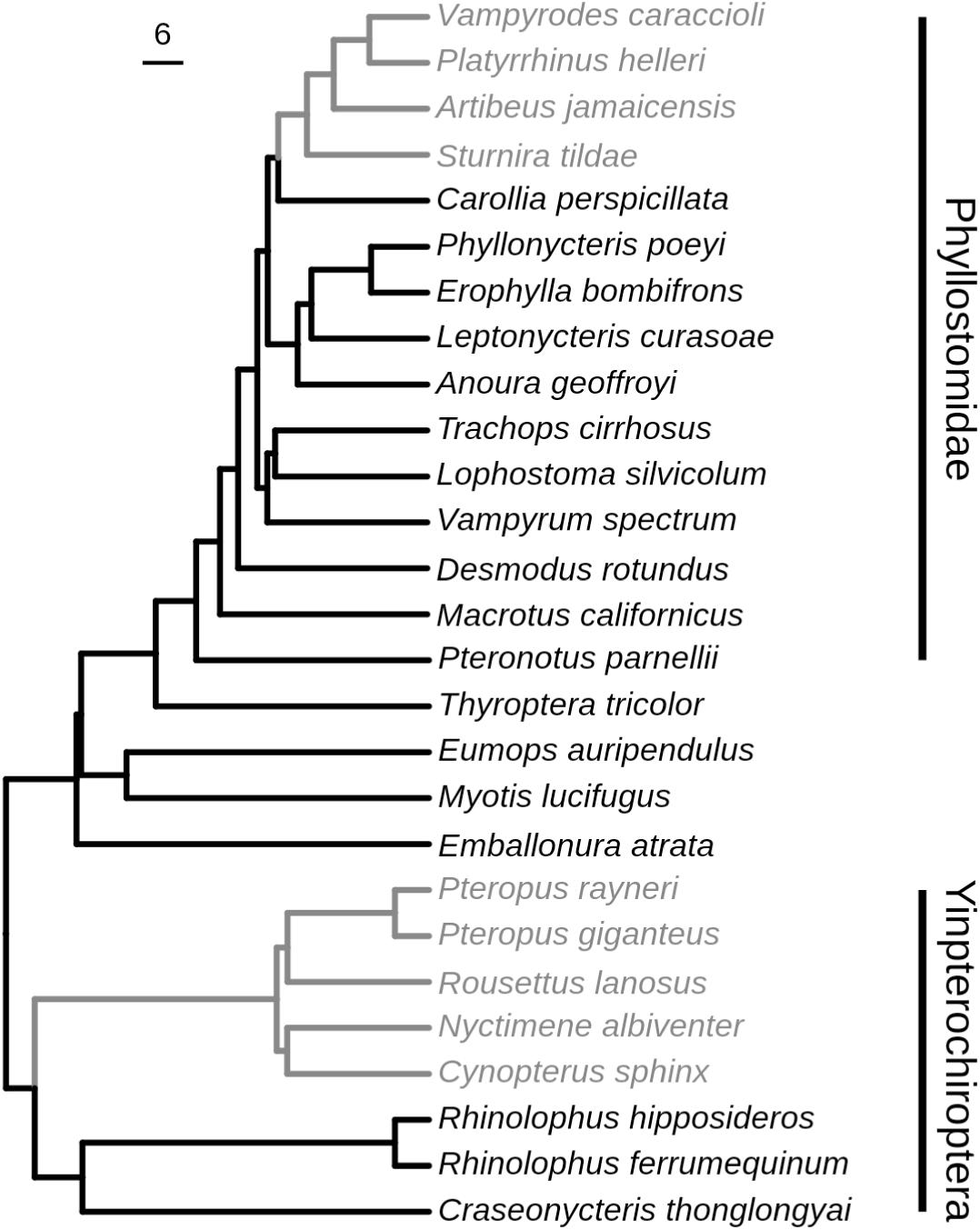
Bat species and clades used in the analysis of topology. Species with branches highlighted in grey are frugivorous.

We use the *OR* gene family in bats, the second most diverse mammalian order with diverse ecological specializations previously linked to *OR* evolution, to test the association between divergent ecologies and olfactory receptors and evaluate model performance. Bats (Mammalia: Chiroptera) are an ideal system in which to test this - while most bats are insectivorous and strongly rely on echolocation to find food resources, several lineages of bats have independently evolved to feed on plants (Jones et al. 2005). Behavioral studies suggest they use scents to find fruiting trees (Korine and Kalko 2005). Previous analyses of the bat olfactory subgenome related specific gene subfamilies to the evolution of frugivory (Hayden et al. 2014), however tree heterogeneity was unexplored as a source of error. Since then, heterogeneity in diversification across the entire bat phylogeny was traced to a single branch that also corresponds to a shift toward a primarily frugivorous diet (Shi and Rabosky 2015), and misleading effects of this heterogeneity on -SSE analyses have been described (Rojas et al. 2018).

Given the potential confounding effects of tree heterogeneity on birth-death equilibrium, we used simulations to determine if such an effect was present in models integrating the birth-death process with the phylogeny. We also used two approaches, both based on a combination of gene and species trees, to infer adaptation in olfactory receptors to the new frugivorous diet: Poisson mixed models that estimate rates of birth and death (Sackton et al. 2017), and phylogenetic instability analyses that uses gene-tree/species-tree reconciliation (Curran et al. 2018). We hypothesize that one or more *OR* gene families involved in shifts to frugivory have expanded in number relative to other *OR* gene families across bats. Changes linked to frugivory could then be identified as a shift in duplication or loss rate (increased duplication or decreased loss in frugivorous vs. non-frugivorous bats), or unusual discordance between these gene trees and the species tree. The results confirmed tree heterogeneity as a confounding factor in testing multi-gene adaptation, identified gene subfamilies experiencing higher turnover, and uncovered high discordance linked to the evolution of frugivory.

## Results

### Influence of topology on CAFE results

We randomly permuted observed olfactory receptor (*OR*) gene copy numbers on inferred, non-Yule species trees of Yangochiroptera and Yinpterochiroptera, comparing single-λ (single-rate) and two-λ (two-rate) models. This revealed that CAFE has a false positive rate on the yangochiropteran tree between 26% and 69%, and between 0% and 76% on the yinpterochiropteran tree (table 1). When branch lengths were Yule-transformed, the false positive rates for the yangochiropteran tree increased for all but one OR gene subfamily (gene subfamily 4) to between 44% and 91%. The false positive rates for the Yule-transformed yinpterochiropteran tree decreased to 0% - 0.1%.

**Table 1.** Proportion of false positives by tree and gene subfamily.

| Gene | Yangochiroptera | Yinpterochiroptera | Yule-transformed Yangochiroptera | Yule-transformed Yinpterochiroptera |
| --- | --- | --- | --- | --- |
| 1/3/7 | 0.36 | 0 | 0.88 | 0 |
| 2/13 | 0.28 | 0.37 | 0.55 | 0 |
| 4 | 0.46 | 0.74 | 0.44 | 0.001 |
| 5/8/9 | 0.26 | 0.22 | 0.86 | 0 |
| 6 | 0.57 | 0.70 | 0.90 | 0 |
| 11 | 0.69 | 0.74 | 0.91 | 0 |
| 52 | 0.65 | 0.76 | 0.57 | 0 |

### Duplication and Loss Events (Notung)

We used Notung to estimate the number of gene duplication and loss events for each OR gene subfamily. The number of duplication events inferred per OR gene subfamily per species or internal branch ranged from 0 to 44 (maximum in OR gene subfamily 1/3/7 in *Anoura geoffroyi*), and the number of loss events ranged from 0 to 90 (maximum in OR gene subfamily 5/8/9 in *Desmodus rotundus*). Most branches had relatively more loss events than duplication events, implying a decrease in gene family size (figure 3).

**Figure 3.**
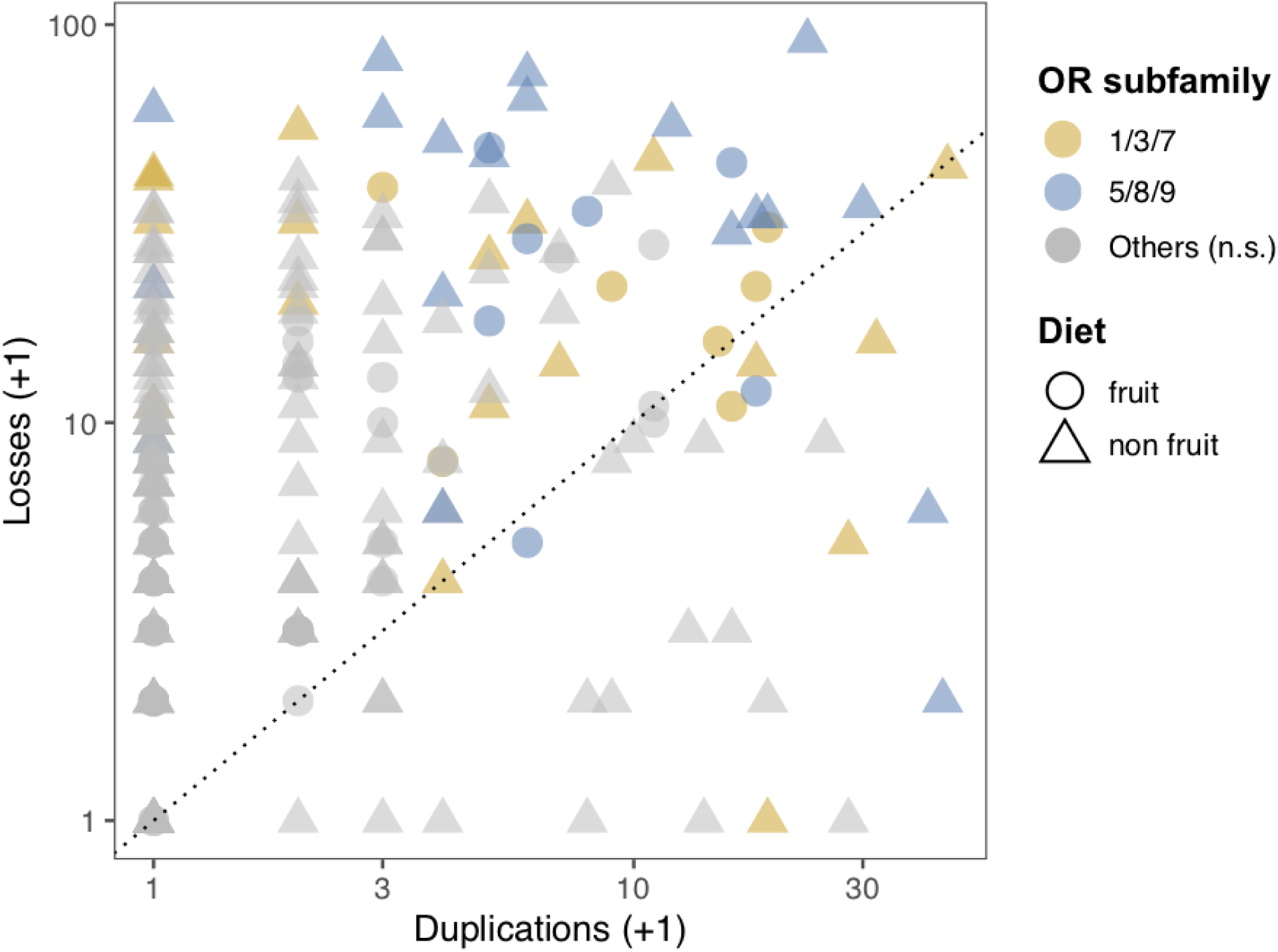
The number of loss events vs the number of duplication events for each olfactory receptor gene subfamily in each phyllostomid species. Triangles represent non-frugivorous species (animalivores and omnivores), circles represent frugivorous species. Colored dots correspond to *OR* gene families in which the number of duplication events was found to be significantly different between frugivorous and non-frugivorous species.

### Poisson overdispersion model

Two *OR* gene subfamilies were identified as having more duplication events based on their posterior predictive intervals excluding 0, *OR* 1/3/7 and *OR* 5/8/9. *OR* 1/3/7 was associated with a mean 1.26X increase in duplication events, and *OR* 5/8/9 was associated with a mean 1.34X increase in duplication events. However, diet did not predict duplication events (figure 4). There is some evidence for increases in duplication events in two combinations of diet and gene subfamily: While the 90% credible intervals of *OR* 2/13, *OR* 6, and the interactions of *OR* 1/3/7 and *OR* 5/8/9 with frugivory included 0, their 70% credible intervals did not (figure 4).

**Figure 4.**
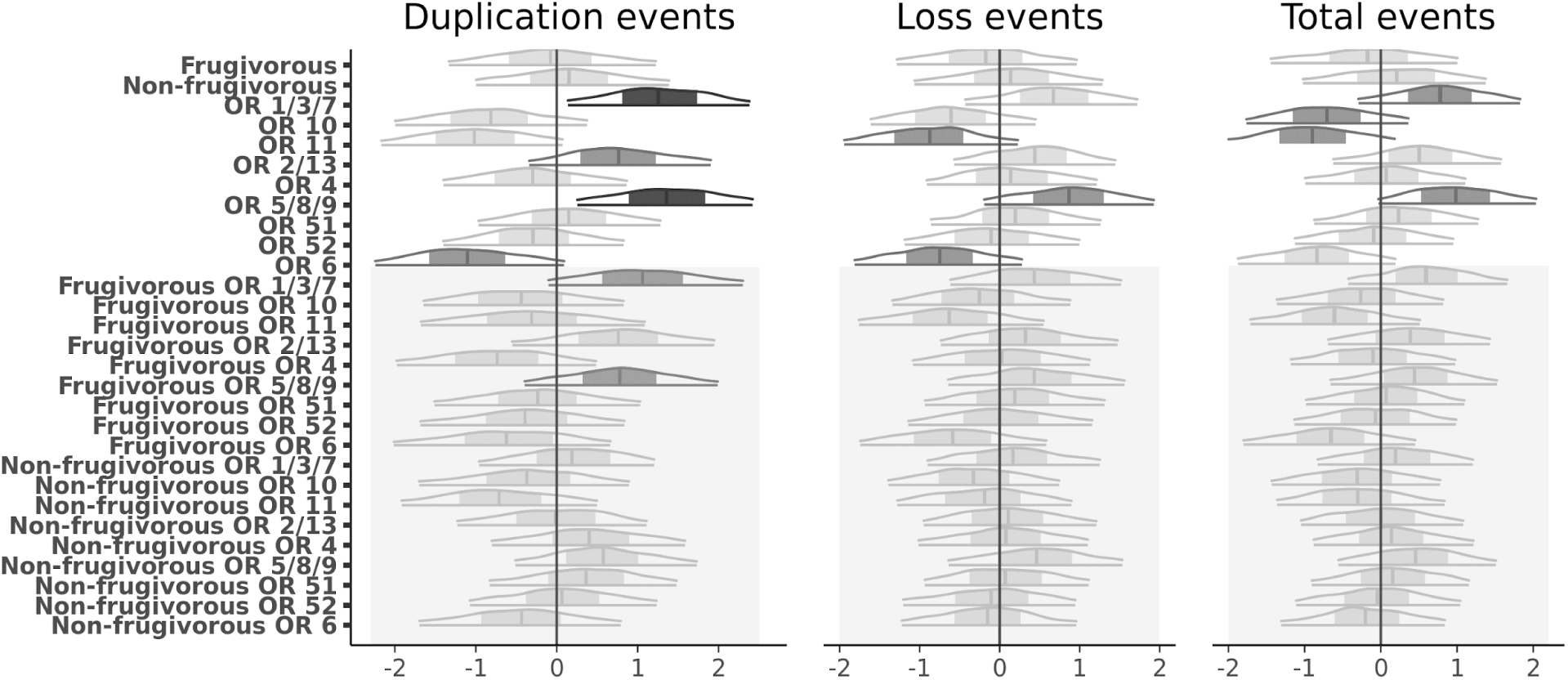
The effect of diet (frugivorous vs non-frugivorous), *OR* gene family, and *OR* gene family in only frugivorous or non-frugivorous species on the number of duplication events (first panel), the number of loss events (second panel), and the total number of duplication and loss events per branch (third panel). Distributions outline the 90% credible intervals of the coefficient of the effect of each predictor on the total number of duplication events per branch, with the 50% credible intervals shaded. Distributions in dark grey have 90% credible intervals that do not include 0, distributions in medium grey have 70% credible intervals that do not include 0, and distributions in light grey have 70% credible intervals that do include 0. Grey boxes highlight the diet x *OR* family interaction terms.

None of the three sets of predictors tested (*OR* gene family, diet, and the interactions of *OR* gene family and diet) predicted either loss events, or the total number of duplication/loss events per species based on the 90% credible intervals. However, at 70% credible intervals, *OR* 11 and *OR* 6 had fewer loss events, and *OR* 5/8/9 more loss events (figure 4). *OR* 10 and *OR* 11 had fewer total duplication/loss events, and *OR* 1/3/7 and *OR* 5/8/9 had more total duplication/loss events (figure 4).

### Phylogenetic instability

Three *OR* gene families, *OR* 5/8/9, *OR* 2/13, and *OR* 6, had significant differences in instability scores between frugivores and non-frugivores based on a Mann-Whitney U test (*OR* 5/8/9: Z = 3.77, p = 0.000165; *OR* 2/13: Z = 7.36, p = 1.84 * 10^−13^; *OR* 6: Z = - 2.47, p = 0.0135) before correcting for multiple comparisons, but only two (*OR* 5/8/9 and *OR* 2/13) were significant after a Hommel correction (figure 5). Frugivorous bats had significantly higher rank instability scores than non-frugivorous bats for both *OR* 5/8/9 and *OR* 2/13. While *OR* 6 was not significant after correcting for multiple comparisons, there was a trend toward frugivorous bats having lower instability scores than non-frugivorous bats (Sup. figure 1A).

**Figure 5.**
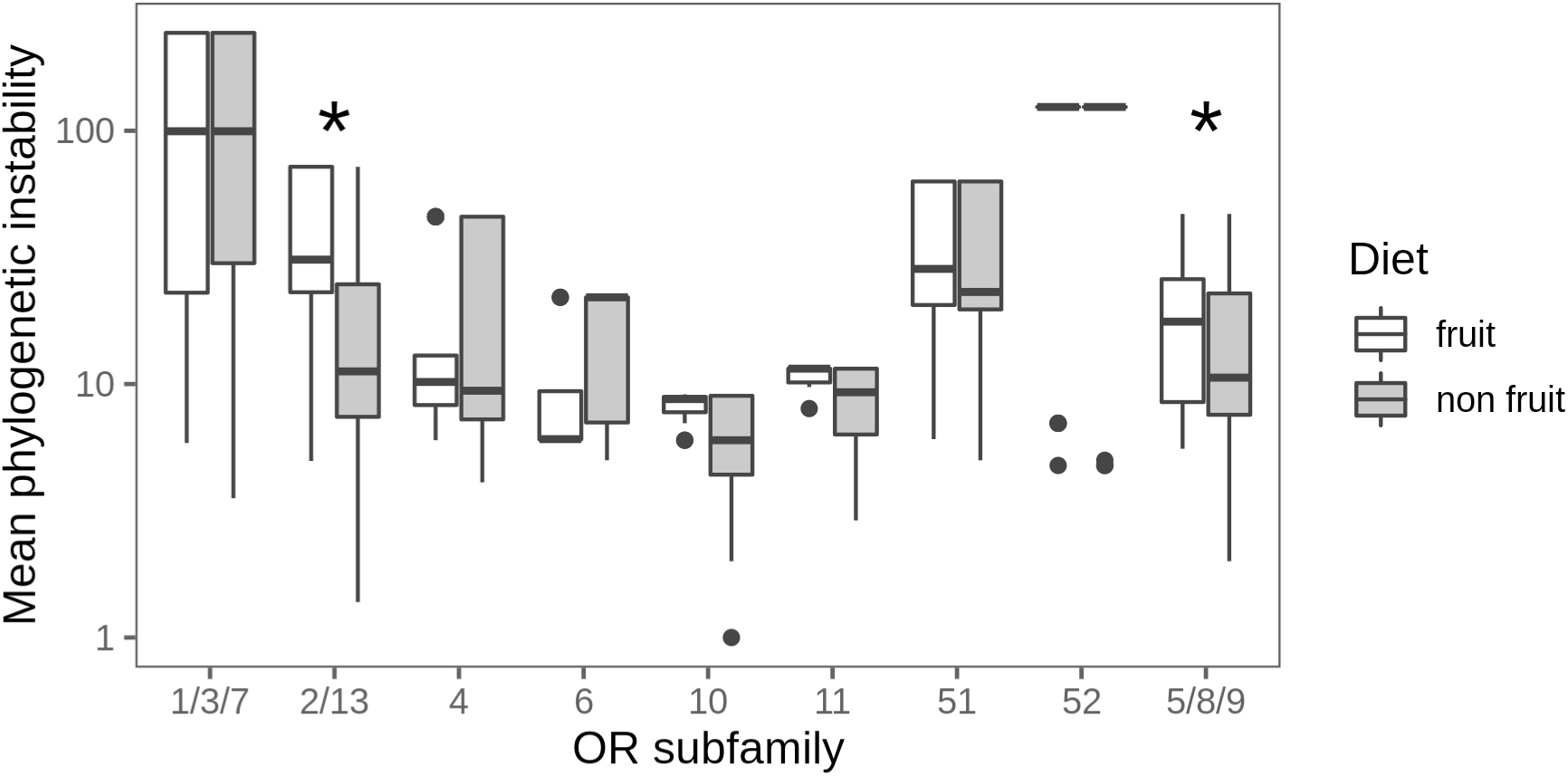
Average instability scores of frugivorous (white) vs. non-frugivorous (grey) bat species in each *OR* gene family. Asterisks designate *OR* subfamilies with significant differences in rank instability scores between frugivorous and non-frugivorous species.

*OR* 5/8/9 and *OR* 2/13 had significant differences in instability between frugivorous and non-frugivorous species, and were the two *OR* subfamilies with the most orthologs (Sup. figure 1B, C). *OR* 5/8/9 contained 580 genes and was sorted into 80 groups, with instability scores ranging from 1.00 to 46.97. *OR* 2/13 contained 239 genes and was sorted into 27 instability groups, with scores ranging from 1.38 to 72.18. *OR* 6, however, was a relatively smaller subfamily, containing only 35 genes which were sorted into 7 instability groups. *OR* 10 is an example of a subfamily with little variation between diets. There are fewer orthologs in the subfamily, and each instability group only had between one and five genes in it (Sup. figure 1D).

## Discussion

We show that species tree heterogeneity misleads inference of gene duplication and loss rates, with implications for tests of ecological associations (e.g., Chang and Duda 2012; Dahan et al. 2015; Ramasamy et al. 2016). Although the influence of tree heterogeneity on analyses of speciation rates has been demonstrated (Rabosky and Goldberg 2015; Rojas et al. 2018), the conflation of tree heterogeneity with rate shifts in multi-gene family evolution had not been previously demonstrated. We show that the most commonly used method for modeling selection on gene copy number, CAFE (Bie et al. 2006; Han et al. 2013), is prone to error with unbalanced species trees, with implications for analyses of many empirical data sets. This type of error, however, can be overcome by adopting tree transformations, or using alternative methods, as shown here. Therefore, whenever tree heterogeneity is suspected, simulations such as those presented here can help determine what methods are appropriate for testing links between rates of multi-gene family evolution and specific clades or branches.

Using simulations we find CAFE has an unacceptably high false positive rate in non-Yule trees, making it inappropriate for analyses of many empirical systems. CAFE provides an elegant model comparison framework for testing for duplication (and/or loss) rate shifts at specific hypothesized nodes of a species phylogeny, and is widely applied for this purpose across many systems. However, its framework assumes a Yule tree, a premise rarely tested for empirical phylogenies. The high false positive rate we show is the result of heterogeneity in the tree shape (non-Yule branching patterns and branch lengths) being ascribed to a duplication or loss rate shift. In effect, CAFE is detecting a deviation from a neutral, single-rate model, but in a non-Yule tree that deviation may come from tree shape instead of from rate shifts in gene family evolution.

While the spurious association between a non-null model and a factor being tested (in this case, duplication/loss rate shifts) was first demonstrated with binary state-speciation and extinction models (BiSSE) (Rabosky and Goldberg 2015), this challenge is a more general consequence of the integration of the species phylogeny with rates superimposed therein, and has also been found with continuous data (Harvey and Rabosky 2018; Rojas et al. 2018). Solutions and alternatives have been formulated for SSE-based models (e.g., (Beaulieu and O’Meara 2016; Rabosky and Huang 2016; Harvey and Rabosky 2018; May and Moore 2019), but are unavailable for this newly-identified instance of tree imbalance producing spurious results. Our simulations show transforming the species tree to meet the Yule assumption can overcome the conflation of heterogeneity in the species tree with gene family rate shifts. This, however, must be tested for particular cases. Methods that do not make assumptions about the tree shape are available as alternatives.

In an empirical application, and using methods that are not confounded by the effects of tree heterogeneity on birth-death equilibrium, we show that despite an overall decrease in most olfactory receptor (*OR*) gene subfamily sizes across phyllostomid bat species, certain *OR* subfamilies have expansions associated with a dietary shift to frugivory. These results contradict some of the findings of a previous study associating *OR* subfamily size changes with a shift to frugivory in phyllostomid bats (Hayden et al. 2014), suggesting that collapsing the gene subfamily data into principal components allows tree-wide instability in a gene subfamily to be misinterpreted as having ecological associations. Our analyses overcome this shortcoming by using reconciliation to analyze rate shifts directly, revealing in greater depth in the dynamics of duplication and loss events associated with frugivory.

By using alternative methods not dependent on tree topology, we identified an increase in duplication events in *OR* subfamily 5/8/9 as linked with the shift to frugivory in phyllostomid bats. Previous work also found a significant increase in duplication events in this subfamily (Hayden et al. 2014), but for non-frugivorous phyllostomids instead. We explain this contradiction through the interaction between duplication events and methods of analysis. The number of duplication events in *OR* subfamily 5/8/9 suggests expansion across phyllostomids, frugivorous and non-frugivorous (figure 3 this paper, Hayden et al. figure 4). Using methods that directly examine the interaction of the number of duplication events and dietary category (Poisson overdisperson) and the deviation of duplication patterns according to diet (phylogenetic instability), we were able to measure the association of duplication events and diet despite an overall increase in the number of duplication events in *OR* subfamily 5/8/9 across the tree.

Analyses by Hayden et al. 2014 added even more abstraction by collapsing *OR* subfamily variation into principal components. This approach compressed the data in ways that could prove consequential. First, while the principal components can relate back to specific gene subfamilies, variation across multiple subfamilies influences their value, making it difficult to discover subfamily correlates of ecology. Second, there was no explicit model of ecological trait evolution and comparisons of the principal components using ANOVA were limited to the tips and secondarily inferred to be localized to internal branches. This highlights the importance of methods that use the number of duplication events directly, as is evident in the resulting instability in the *OR* subfamily 5/8/9 gene tree (Sup. figure 1C).

Based on its expansion within frugivorous phyllostomids, elements of *OR* subfamily 5/8/9 may therefore specifically respond to volatile organic compounds (VOCs) released by ripening fruits. Indeed, expansions of this subfamily have recently been linked to herbivory in mammals (Hughes et al. 2018), lending confidence to the conclusion that *OR* subfamily 5/8/9 is involved in detecting plant VOCs. It has been previously hypothesized that the size of an *OR* repertoire could influence olfactory sensitivity {\rtf (Rouquier et al. 2000; but see Wackermannová et al., 2016), and some evidence suggests this may be the case (Laska and Shepherd 2007; Rizvanovic et al. 2013). These additional copies of subfamily 5/8/9 genes could increase a bat’s perception of fruit VOCs. However, this could be either because of increased sensitivity to a single VOC as a result of having more receptors for that specific VOC, or increased discrimination between many similar VOCs, resulting from slight functional differences between *OR* gene copies within the subfamily (Laska and Galizia 2001; Rizvanovic et al. 2013). To discern whether sensitivity or discrimination is responsible for this expansion, individual-level differences in *OR* gene subfamily repertoires and associated VOC preferences should be integrated with models of gene subfamily birth/death dynamics.

OR subfamily 2/13 was also previously found to be important in the shift to frugivory within phyllostomids (Hayden et al. 2014), and this is corroborated by phylogenetic instability scores in this study (figure 5). In contrast, the number of duplication events within this subfamily was not higher in frugivorous species, as determined by posterior predictive intervals (figure 4). While related, the number of duplication events and phylogenetic instability cannot be expected to always yield consistent results. Incongruence (instability) could increase at a branch without quite pushing the number of events at the ancestral branch toward significance. Alternatively, an expansion in this subfamily could be associated with individual branches only partially coincident with a shift to frugivory, instead of the single branch associated with frugivory itself.

While *OR* subfamily 1/3/7 was previously found to be important in the shift to frugivory within phyllostomids (Hayden et al. 2014), our results suggest this subfamily is highly unstable across the whole phyllostomid tree. Within fruit-specialized phyllostomids, there is a residual trend toward more duplications, but this subfamily experiences high turnover across the whole tree. Thus, its expansion may be related to either an inherent genetic mechanism, or an ecological factor affecting all of Phyllostomidae, rather than the shift to frugivory within the family.

We have shown that expansions of at least one *OR* subfamily, 5/8/9, are associated with a shift to frugivory in phyllostomid bats. Nonetheless, we have also shown that expansions in *OR* subfamilies are not always associated with an important ecological change. For example, the results of the Poisson overdispersion model show that about half of the subfamilies tested may simply have more duplication events than expected based on the phylogeny, regardless of diet. This means other factors can confound these analyses, some of which will often be unaccounted for. By combining a method that measures variation in the number of duplication(/loss) events (Poisson overdispersion model) with one that measures deviation from the expected number of copies (phylogenetic instability), we are able to identify *OR* subfamilies for which conflicting results (e.g., *OR* subfamily 2/13) point to confounding factor(s). Changes in the per-lineage numbers of duplication and loss events not accounted for in the shift to frugivory may also be driven by other behavioral and ecological factors (e.g., mate identification, habitat type), or changes in the rates of duplication events for specific genomic regions.

Within *OR* subfamily 5/8/9, and in other *OR* subfamilies found to be associated with ecological shifts in other systems, it is now important to identify whether the subfamily expansion is beneficial because of increased sensitivity or increased discrimination. That is, do more copies of an *OR* gene increase the likelihood that the animal will detect a scent when there are few molecules, or increase the likelihood that the animal will be able to sort out a particular scent of interest from many similar ones? Determining functional differences between *OR* genes within a subfamily, by comparing amino acid substitutions with different physiochemical properties, or by *in vivo* or *in vitro* response experiments, will be the next step to making this distinction.

Testing the adaptive significance of gene subfamily size or changes in gene duplication or loss rates is complex and potentially confounded by the relationship between the birth-death process of gene subfamily evolution and the species phylogeny. This study confirms that, as with state-dependent speciation models, species tree heterogeneity is a confounding factor in inferring multi-gene adaptation, and can cause spurious associations between shifts in birth-death rate and species ecology. Since testing the ecological associations of gene subfamily size and/or duplication and loss rates is a necessary step in determining their adaptive significance, it is thus imperative to use methods that are not confounded by the effects of tree heterogeneity on birth-death equilibrium.

## Methods

### Sequence data and copy number

Aligned consensus sequences of bat *OR* gene families and copy number counts for nine *OR* gene subfamilies (*OR* 1/3/7, *OR* 2/13, *OR* 4, *OR* 5/8/9, *OR* 6, *OR* 10, *OR* 11, *OR* 51, and *OR* 52) in 14 species of Phyllostomids and eight species of Yinpterochiropterans were obtained from Hayden et al. (2014) (figure 2).

### Influence of topology

In order to test the variation in duplication rates in gene families, CAFE assumes a birth-death (Yule) species tree (Han et al. 2013). Many species trees, however, do not fit this assumption. To test the performance on real phylogenetic trees, we ascertained the false positive rate of changes in the duplication rate λ associated with a shift to frugivory in bats. We used both actual species trees and those trees transformed to meet Yule branching time expectations, using copy number counts from Hayden et al. (2014). We ran CAFE independently for *OR* gene families in two bat clades, family Phyllostomidae (poor fit to Yule expectations) and suborder Yinpterochiroptera (better fit to Yule expectations) (figure 2).

For each gene subfamily and species tree, copy number counts from Hayden et al. (2014) were randomized across the tips in 100 permutations, and CAFE was run on each permutation. This randomization preserved observed magnitudes and relative copy number counts, but associated each with a different tip, thus removing the influence and signal of selection on copy numbers corresponding to ecological traits.

Each set of CAFE runs included both a single-λ and a two-λ model, with the shift in λ corresponding to the shift to frugivory. Models were compared via likelihood ratio test using CAFE’s built-in *lhtest* command, which creates a null distribution of likelihood ratios for the null (single-λ) hypothesis. Any significant likelihood ratio test, indicating a better fit to the data for the two-λ model, was thus a false positive for selection. A false positive implies spurious influence of tree shape on the CAFE test, especially if false positives are more prevalent in trees with a poorer fit to a Yule model.

To further discern the influence of non-Yule branch length vs. topology on the false positive rate, we transformed branch lengths of both trees to meet Yule expectations by fitting a Yule model to each tree with yule(), applying Yule expected branching times simulated to fit that model with sim.bdtree(), and applying those branching times to each original tree with compute.brtime() from the ape (Paradis et al. 2004) and geiger (Harmon et al. 2008) packages in R (R Core Team 2018).

### Inferring Gene Trees and Estimating Duplication and Loss Events

We used ModelOMatic (Whelan et al. 2015) to determine the best substitution model for each *OR* gene subfamily, and used garli (Zwickl 2006) to infer gene trees from the alignments provided by Hayden et al. (2014). We then ran Notung (Chen et al. 2000) with default duplication (1.5) and loss (1.0) costs to reconcile the resulting gene and species trees and obtain estimates of the number and location of duplication and loss events.

### Poisson overdispersion model

To determine the influence of diet on the duplication and loss rates of each *OR* gene family, we built a Poisson generalized mixed model of diet, *OR* gene subfamily, and the interaction effect between diet and *OR* gene subfamily, based on the model from Sackton et al. (2017). Although the Sackton et al. (2017) implementation used a maximum likelihood implementation of hierarchical mixed Poisson regressions, here we apply Bayesian method to account for overdispersion in the counts. The model was run using MCMCglmm in the MCMCglmm package (Hadfield 2010) including a branch length offset as a fixed effect to control for species relationships, and all other characters as random effects after Gelman and Hill (2006).

We ran three separate models, one with the number of duplication events per branch and tip, one with the number of loss events, and one with the total number of events as the response variable. The bayesplot package (Gabry et al. 2019) was used to visualize posterior probability distributions.

### Phylogenetic instability

We determined phylogenetic instability for all nine *OR* gene subfamilies in Phyllostomids (figure 2) using MIPhy (Minimizing Instability in Phylogenetics) (Curran et al. 2018) and gene and species trees from Hayden et al. (2014). MIPhy uses patterns of duplications and losses in gene trees to identify groups of genes that are under selection based on deviations from expected baseline patterns. MIPhy quantifies instability in each gene, and assigns each group a score based on how its pattern of gene events differs from what would be expected if the relationships in the gene tree mirrored those of the species tree. A higher instability score indicates that the gene tree is discordant with the species tree, and suggests that it is under positive or negative selection.

To quantify the phylogenetic instability for each of the nine *OR* gene families, we used the MIPhy online web tool with default weights (Curran et al. 2018). The species tree and the list of *OR* genes corresponding to each species were compiled into an information file compatible with MIPhy.

To determine if the resulting instability scores were more likely to be associated with frugivorous species than expected, we performed a 2-tailed Wilcoxon-Mann-Whitney test with species classified as frugivorous or non-frugivorous, using the wilcoxon_test function in the coin packages to account for ties (Hothorn et al. 2019 Aug). No correction for species relatedness was performed because the instability scores already take this into account. A greater score of the genes in an *OR* subfamily would suggest that the family was under selective pressure during the transition to frugivory, causing it to deviate from the expected pattern of duplications and losses based on the gene tree. We applied the Hommel correction for multiple comparisons.

## Acknowledgements

We thank Katie Martin and Sharlene Santana for comments and feedback on the work. The authors would like to thank Stony Brook Research Computing and Cyberinfrastructure, and the Institute for Advanced Computational Science at Stony Brook University for access to the high-performance SeaWulf computing system, which was made possible by a $1.4M National Science Foundation grant (#1531492). This work was supported in part by the National Science Foundation (DEB 1442142, DEB 1456455, and DEB 1838273 to L.M.D., DEB 1701414 to L.M.D. and L.R.Y., PRFB 181203 to L.R.Y.), and the Tinker Foundation and American Association of University Women fellowships to M.E.L.

## References

Assis R, Bachtrog D. 2013. Neofunctionalization of young duplicate genes in Drosophila. Proc Natl Acad Sci. 110(43):17409–17414. doi: 10.1073/pnas.1313759110.

Beaulieu JM, O’Meara BC. 2016. Detecting Hidden Diversification Shifts in Models of Trait-Dependent Speciation and Extinction. Syst Biol. 65(4):583–601. doi: 10.1093/sysbio/syw022.

Bie TD, Cristianini N, Demuth JP, Hahn MW. 2006. CAFE: a computational tool for the study of gene family evolution. 22(10):1269–1271. doi: 10.1093/bioinformatics/btl097.

Chang D, Duda TF. 2012. Extensive and Continuous Duplication Facilitates Rapid Evolution and Diversification of Gene Families. Mol Biol Evol. 29(8):2019–2029. doi: 10.1093/molbev/mss068.

Chen K, Durand D, Farach-Colton M. 2000. NOTUNG: A Program for Dating Gene Duplications and Optimizing Gene Family Trees. J Comput Biol. 7(3–4):429–447. doi: 10.1089/106652700750050871.

Curran DM, Gilleard JS, Wasmuth JD. 2018. MIPhy: identify and quantify rapidly evolving members of large gene families. PeerJ. 6:e4873. doi: 10.7717/peerj.4873.

Dahan RA, Duncan RP, Wilson AC, Dávalos LM. 2015. Amino acid transporter expansions associated with the evolution of obligate endosymbiosis in sap-feeding insects (Hemiptera: sternorrhyncha). BMC Evol Biol. 15(1):52. doi: 10.1186/s12862-015-0315-3.

Gabry J, Simpson D, Vehtari A, Betancourt M, Gelman A. 2019. Visualization in Bayesian workflow. J R Stat Soc Ser A Stat Soc. 182(2):389–402. doi: 10.1111/rssa.12378.

Gelman A, Hill J. 2006. Data Analysis Using Regression and Multilevel/Hierarchical Models. Cambridge University Press (Analytical Methods for Social Research).

Hadfield JD. 2010. MCMC Methods for Multi-Response Generalized Linear Mixed Models: The MCMCglmm *R* Package. J Stat Softw. 33(2). doi: 10.18637/jss.v033.i02. [accessed 2019 Sep 7]. http://www.jstatsoft.org/v33/i02/.

Han MV, Thomas GWC, Lugo-Martinez J, Hahn MW. 2013. Estimating gene gain and loss rates in the presence of error in genome assembly and annotation using CAFE 3. Mol Biol Evol. 30(8):1987–97. doi: 10.1093/molbev/mst100.

Harmon LJ, Weir JT, Brock CD, Glor RE, Challenger W. 2008. GEIGER: investigating evolutionary radiations. Bioinformatics. 24(1):129–131. doi: 10.1093/bioinformatics/btm538.

Harvey MG, Rabosky DL. 2018. Continuous traits and speciation rates: Alternatives to state-dependent diversification models. Cooper N, editor. Methods Ecol Evol. 9(4):984–993. doi: 10.1111/2041-210X.12949.

Hayden S, Bekaert M, Goodbla A, Murphy WJ, Dávalos LM, Teeling EC. 2014. A Cluster of Olfactory Receptor Genes Linked to Frugivory in Bats. Mol Biol Evol. 31(4):917–927. doi: 10.1093/molbev/msu043.

Hothorn T, Winell H, Hornik K, van de Wiel MA, Zeileis A. 2019 Aug. Conditional Inference Procedures in a Permutation Test Framework. CRAN.

Hughes GM, Boston ESM, Finarelli JA, Murphy WJ, Higgins DG, Teeling EC. 2018. The Birth and Death of Olfactory Receptor Gene Families in Mammalian Niche Adaptation. Satta Y, editor. Mol Biol Evol. 35(6):1390–1406. doi: 10.1093/molbev/msy028.

Jones KE, Bininda□Emonds ORP, Gittleman JL. 2005. Bats, Clocks, and Rocks: Diversification Patterns in Chiroptera. Evolution. 59(10):2243–2255. doi: 10.1111/j.0014-3820.2005.tb00932.x.

Laska M, Galizia CG. 2001. Enantioselectivity of Odor Perception in Honeybees (Apis mellifera carnica). Behav Neurosci. 115(3):632–639.

Laska M, Shepherd GM. 2007. Olfactory discrimination ability of CD-1 mice for a large array of enantiomers. Neuroscience. 144(1):295–301. doi: 10.1016/j.neuroscience.2006.08.063.

May MR, Moore BR. 2019. A Bayesian Approach for Inferring the Impact of a Discrete Character on Rates of Continuous-Character Evolution in the Presence of Background-Rate Variation. Syst Biol. syz069.

McGowen MR, Gatesy J, Wildman DE. 2014. Molecular evolution tracks macroevolutionary transitions in Cetacea. Trends Ecol Evol. 29(6):336–346. doi: 10.1016/j.tree.2014.04.001.

Nei M. 1969. Gene Duplication and Nucleotide Substitution in Evolution. Nature. 221(5175):40–42. doi: 10.1038/221040a0.

Nei M, Gu X, Sitnikova T. 1997. Evolution by the birth-and-death process in multigene families of the vertebrate immune system. Proc Natl Acad Sci. 94(15):7799–7806. doi: 10.1073/pnas.94.15.7799.

Nei M, Hughes AL. 1991. Polymorphism and evolution of the major histocompatibility complex loci in mammals. In: Selander R, Clark A, Whittam T, editors. Evolution at the Molecular Level. Sunderland, MA: Sinauer Associates. p. 222–247. [accessed 2019 Dec 5]. http://www.personal.psu.edu/nxm2/1991%20Publications/1991-nei-hughes2.pdf.

Nei M, Rooney AP. 2005. Concerted and Birth-and-Death Evolution of Multigene Families. Annu Rev Genet. 39(1):121–152. doi: 10.1146/annurev.genet.39.073003.112240.

Paradis E, Claude J, Strimmer K. 2004. APE: Analyses of Phylogenetics and Evolution in R language. Bioinformatics. 20(2):289–290. doi: 10.1093/bioinformatics/btg412.

R Core Team. 2018. R: A language and environment for statistical computing. R Found Stat Comput Vienna Austria. https://www.R-project.org.

Rabosky DL, Goldberg EE. 2015. Model Inadequacy and Mistaken Inferences of Trait-Dependent Speciation. Syst Biol. 64(2):340–355. doi: 10.1093/sysbio/syu131.

Rabosky DL, Huang H. 2016. A Robust Semi-Parametric Test for Detecting Trait-Dependent Diversification. Syst Biol. 65(2):181–193. doi: 10.1093/sysbio/syv066.

Ramasamy S, Ometto L, Crava CM, Revadi S, Kaur R, Horner DS, Pisani D, Dekker T, Anfora G, Rota-Stabelli O. 2016. The Evolution of Olfactory Gene Families in *Drosophila* and the Genomic Basis of chemical-Ecological Adaptation in *Drosophila suzukii*. Genome Biol Evol. 8(8):2297–2311. doi: 10.1093/gbe/evw160.

Rizvanovic A, Amundin M, Laska M. 2013. Olfactory Discrimination Ability of Asian Elephants (Elephas maximus) for Structurally Related Odorants. Chem Senses. 38(2):107–118. doi: 10.1093/chemse/bjs097.

Rojas D, Ramos Pereira MJ, Fonseca C, Dávalos LM. 2018. Eating down the food chain: generalism is not an evolutionary dead end for herbivores. Harmon L, editor. Ecol Lett. 21(3):402–410. doi: 10.1111/ele.12911.

Rouquier S, Blancher A, Giorgi D. 2000. The olfactory receptor gene repertoire in primates and mouse: Evidence for reduction of the functional fraction in primates. Proc Natl Acad Sci. 97(6):2870–2874. doi: 10.1073/pnas.040580197.

Sackton TB, Lazzaro BP, Clark AG. 2017. Rapid expansion of immune-related gene families in the house fly, *Musca domestica*. Mol Biol Evol. doi: 10.1093/molbev/msw285. [accessed 2017 Mar 17]. https://academic.oup.com/mbe/article-lookup/doi/10.1093/molbev/msw285.

Vandewege MW, Mangum SF, Gabaldón T, Castoe TA, Ray DA, Hoffmann FG. 2016. Contrasting patterns of evolutionary diversification in the olfactory repertoires of reptile and bird genomes. Genome Biol Evol. doi: 10.1093/gbe/evw013. [accessed 2019 Dec 5]. https://academic.oup.com/gbe/article-lookup/doi/10.1093/gbe/evw013.

Wackermannová M, Pinc L, Jebavý L. 2016. Olfactory Sensitivity in Mammalian Species. Physiol Res. 65:369–390.

Whelan S, Allen JE, Blackburne BP, Talavera D. 2015. ModelOMatic: Fast and Automated Model Selection between RY, Nucleotide, Amino Acid, and Codon Substitution Models. Syst Biol. 64(1):42–55. doi: 10.1093/sysbio/syu062.

Yohe LR, Liu L, Dávalos LM, Liberles DA. 2019. Protocols for the Molecular Evolutionary Analysis of Membrane Protein Gene Duplicates. In: Sikosek T, editor. Computational Methods in Protein Evolution. Vol. 1851. New York, NY: Springer New York. p. 49–62. [accessed 2019 Mar 14]. http://link.springer.com/10.1007/978-1-4939-8736-8_3.

Zwickl DJ. 2006. Genetic algorithm approaches for the phylogenetic analysis of large biological sequence datasets under the maximum likelihood criterion [Dissertation]. The University of Texas at Austin.

