## Supplementary Figure 1 for "Species tree disequilibrium positively misleads models of gene family evolution"

Supplementary Figures

A.

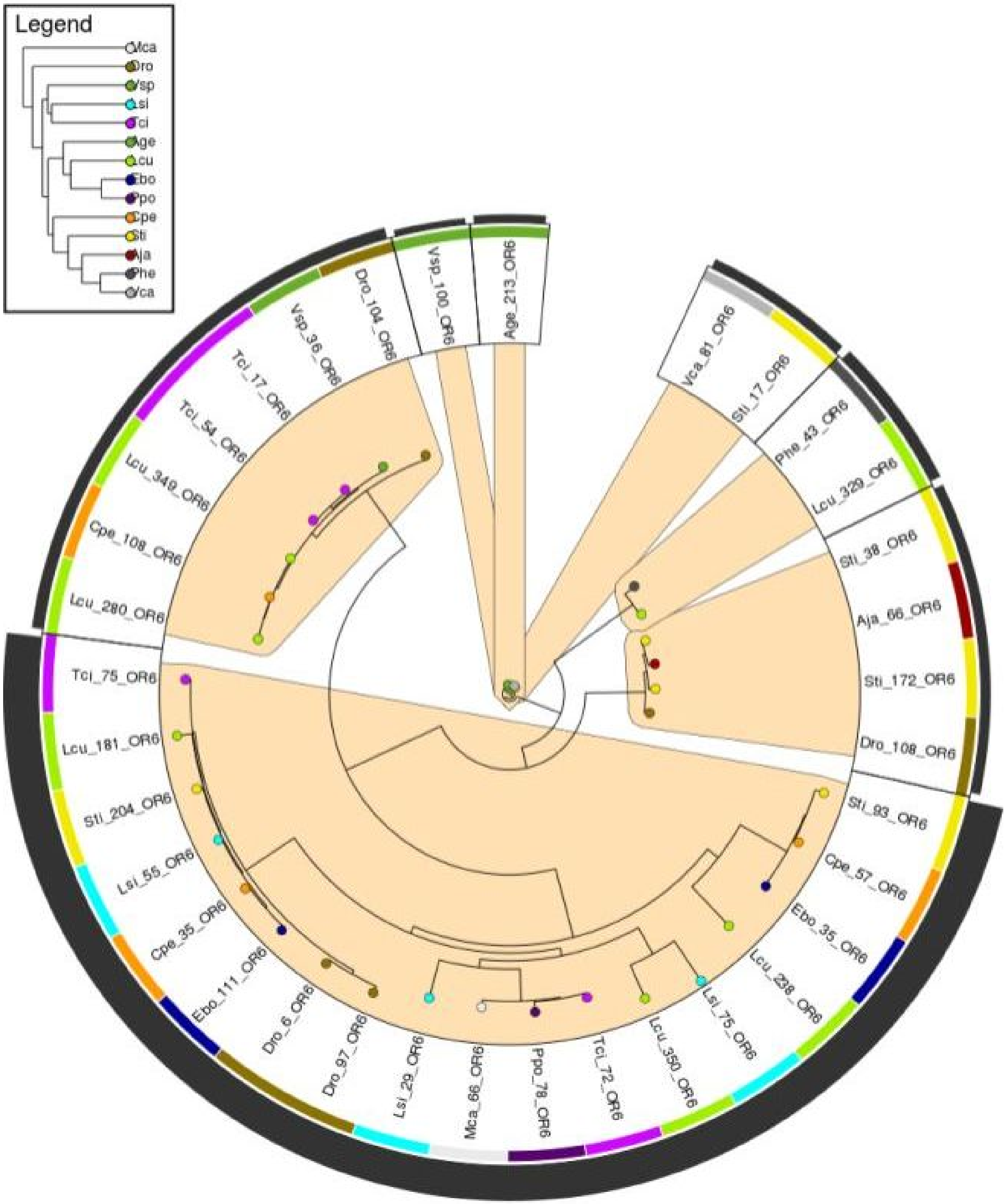

B.

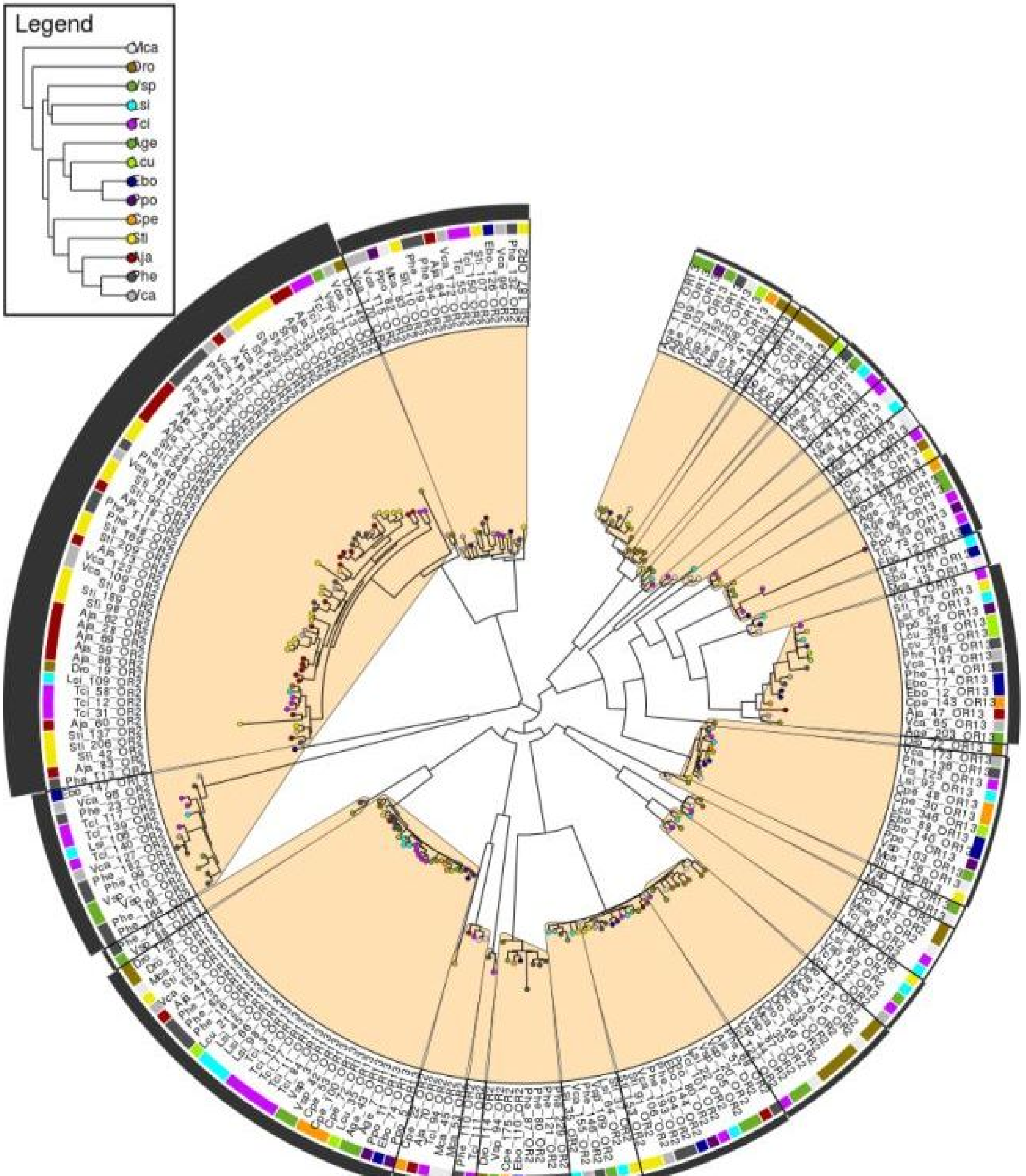

C.

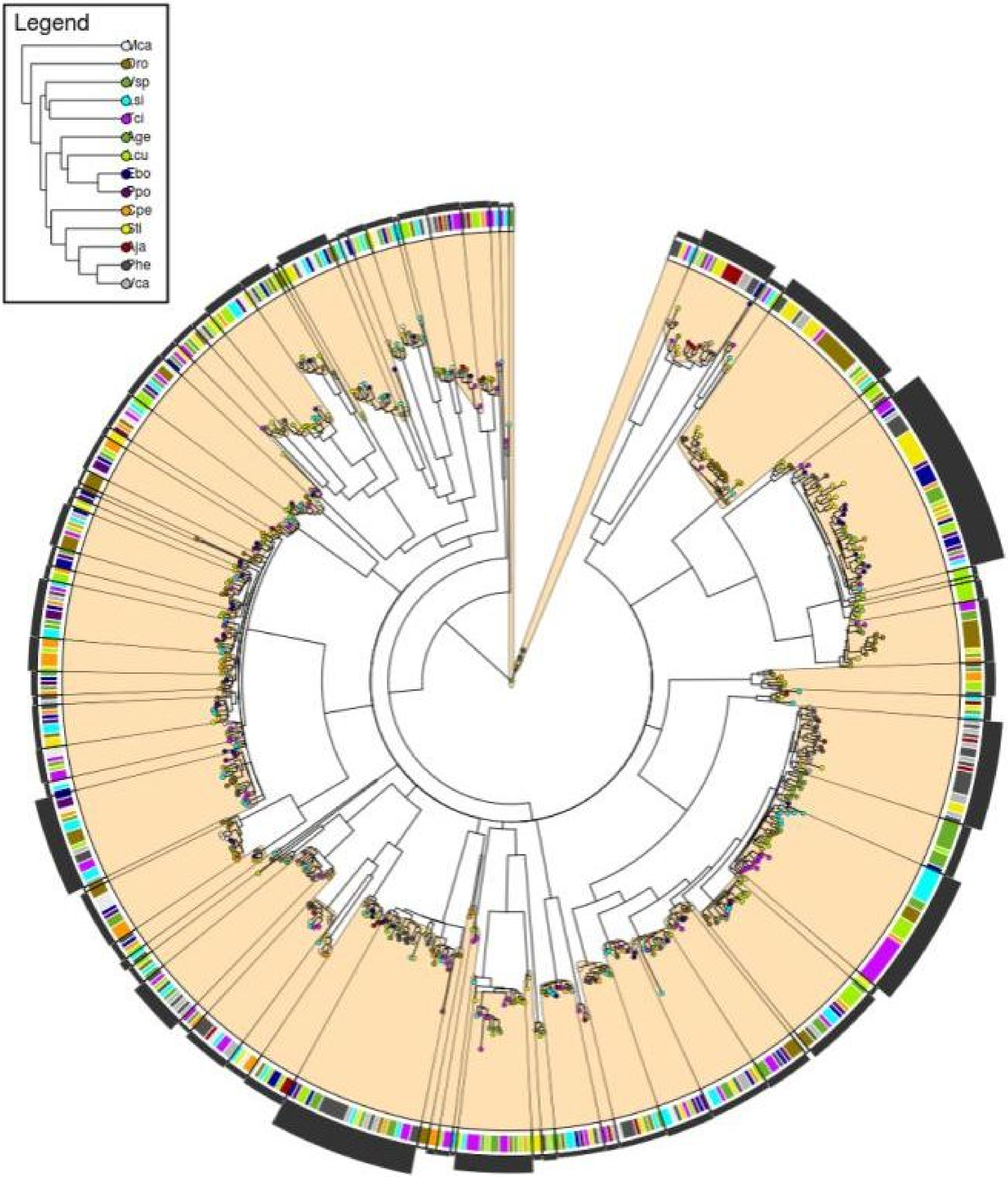

D.

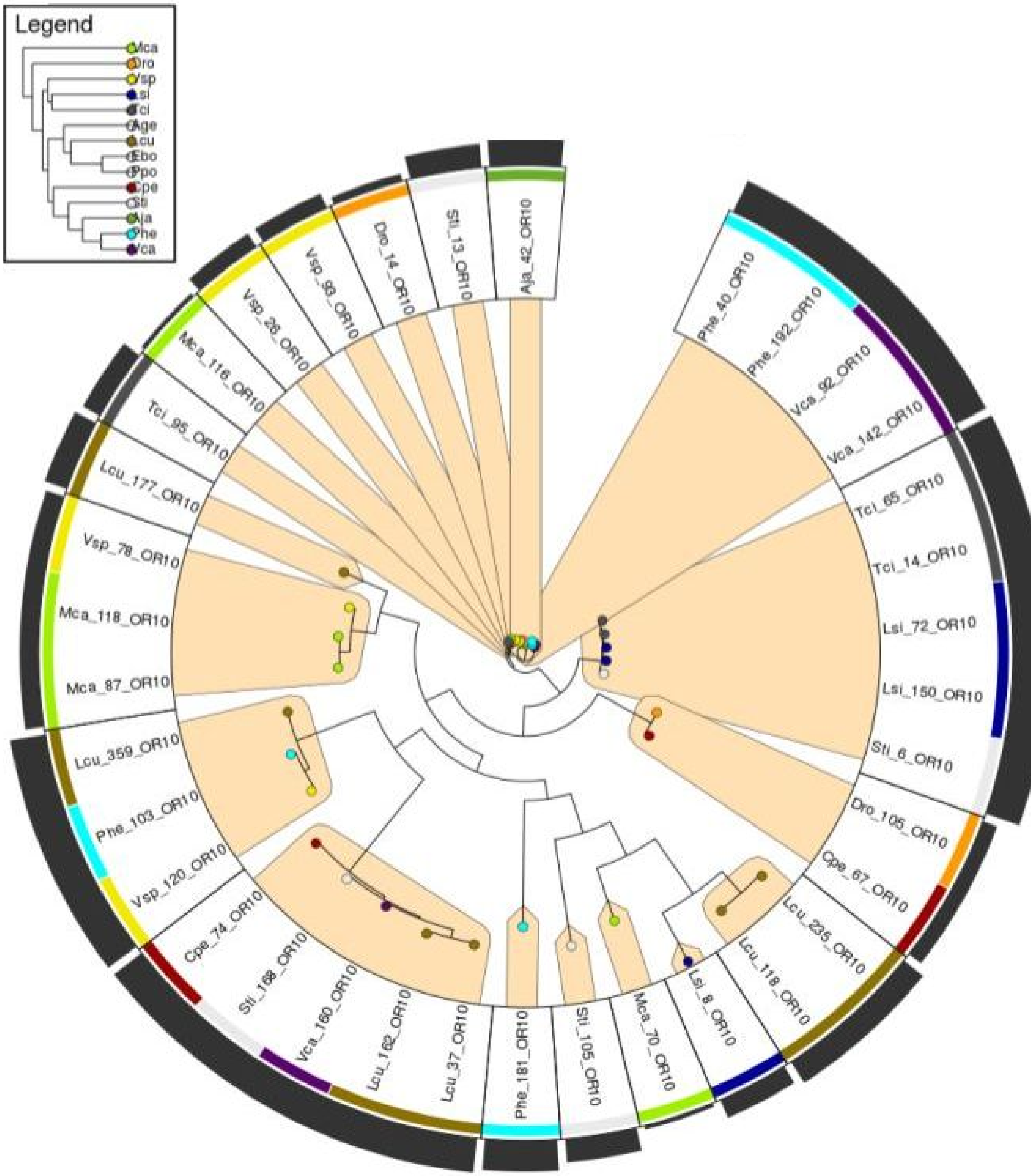
Legend
Mca
Dro
Vsp
Lsi
Tci
Age
Lcu
Ebo
Ppo
Cpe
Sti
Aja
Phe
Vca
Aja_42_OR10
Phe_40_OR10
Phe_192_OR10
Vca_92_OR10
Vca_142_OR10
Tci_65_OR10
Tci_14_OR10
Lsi_72_OR10
Lsi_150_OR10
Sti_6_OR10
Dro_105_OR10
Cpe_67_OR10
Lcu_235_OR10
Lcu_118_OR10
Lsi_8_OR10
Mca_70_OR10
Sti_105_OR10
Phe_181_OR10
Lcu_37_OR10
Lcu_162_OR10
Vca_160_OR10
Sti_168_OR10
Cpe_74_OR10
Vsp_120_OR10
Phe_103_OR10
Lcu_359_OR10
Mca_87_OR10
Mca_118_OR10
Vsp_78_OR10
Lcu_177_OR10
Tci_95_OR10
Mca_116_OR10
Vsp_26_OR10
Vsp_93_OR10
Dro_14_OR10
Sti_13_OR10

Supplementary Figure 1. Gene trees grouped into minimum instability groups in four *OR* gene families. A) *OR* 6, which had a non-significant trend toward lower instability in frugivorous species, B) *OR* 2/13, which had significantly higher instability in frugivorous species, C) *OR* 5/8/9, which had significantly higher instability in frugivorous species, and D) *OR* 10, which had no difference in instability score between frugivorous and non-frugivorous species. Gene tree visualizations are produced by MIPhy, with species identity indicated by color and minimum instability groups highlighted in orange. Insets show the average instability scores for frugivorous (green) and non-frugivorous (grey) species, as in figure 5. Note the different scales of the y-axes between panels.
